# Spatial Congruence Analysis (SCAN): An objective method for detecting biogeographical patterns based on species’ range congruences

**DOI:** 10.1101/2021.01.11.426192

**Authors:** Cassiano A F R Gatto, Mario Cohn-Haft

## Abstract

Similar species ranges may represent outcomes of common biological processes and so form the basis for biogeographical concepts such as areas of endemism and ecoregions. Nevertheless, spatial range congruence is rarely quantified, much less incorporated in bioregionalization methods as an explicit parameter. Furthermore, most available methods suffer from limitations related to the loss, or the excess of range information, or scale bias associated with the use of grids, and the incapacity to recognize independent overlapped patterns or gradients of range distributions. Here, we propose an analytical method, Spatial Congruence Analysis (SCAN), to identify biogeographically meaningful groups of species, called biogeographic elements. Such elements are based on direct and indirect spatial relationships among species’ ranges and vary depending on an explicit measure of range congruence controlled as a numerical parameter in the analysis. A one-layered network connects species (vertices) using pairwise spatial congruence estimates (edges). This network is then analyzed for each species, separately, by an algorithm that accesses the entire web of spatial relationships to the reference species. The method was applied to two datasets: a simulated gradient of ranges and real distributions of birds. The gradient results showed that SCAN can describe gradients of distribution with a high level of detail, without confounding transition zones with true biogeographical units, a frequent pitfall of other methods. The bird dataset showed that only a small portion of range overlaps is biogeographically meaningful, and that there is a large variation in types of patterns that can be found with real distributions. Distinct reference species may converge on similar or identical groups of spatially related species, may lead to recognition of nested species groups, or may even generate similar spatial patterns with no species in common. The biological significance or causal processes of these patterns should be investigated a posteriori. Patterns can vary from simple ones, composed by few highly congruent species, to complex, with numerous alternative component species and spatial configurations, depending on particular parameter settings as determined by the investigator. This approach eliminates or reduces limitations of other methods and permits pattern description without hidden assumptions about processes, and so should make a valuable contribution to the biogeographer’s toolbox.

“If there is any basic unit of biogeography, it is the geographic range of a species.” - Brown, Stevens & Kaufman [1].

“[spatial] congruence […] should be optimized, while realizing that this criterion will most likely never be fully met” - HP Linder [2].

## Introduction

Species ranges are surely determined by physical and biological processes at multiple scales of space and time [1,3]. Although the exact processes and timing are likely to be unique in each case, it is relatively common in nature that several species share the same or similar global distributions [4]. This intriguing phenomenon of shared distributions is an important building block in the study of biogeography, because it may be possible to infer from it the action of shared causes [5,6]. Thus, congruent ranges have long been a major source of insights for naturalists at the foundations of ecology and evolution [7–10]. The concept of congruent ranges is used to this day to describe units in biogeography, such as areas of endemism [11], ecoregions [12], bioregions [13], and chorotypes, among others [4].

Recognition of congruent distributions has traditionally been a relatively subjective process with some unexpressed premises. When species show precisely similar distributions, it is natural to assume that strong ecological pressures play an important role in this congruence. For example, a bird species whose distribution coincides closely with that of a particular tree (e.g., *Picoides tridactyla* and *P. dorsalis* three-toed woodpeckers and *Picea* spp. spruce trees) [14] is readily interpreted as being dependent on the presence of those trees. On the other hand, coincident distributions of dozens of unrelated bird species in discrete parts of the Amazon basin are assumed to reflect past vicariant events that led to differentiation and geographic limitation in numerous groups simultaneously [15,16]. In these latter cases, less precise congruence is implicitly acceptable because each species is likely to have adapted to modern environments in unique and slightly distinct ways.

In an attempt to make spatial relationships more explicit and quantifiable, modern approaches to bioregionalization have adapted methods from ecological [17,18], phylogenetic [19], and network analyses [13], in which sample units (e.g., collecting stations, characters in a phylogenetic matrix) are replaced by grid-cells dividing the studied area. The use of indexes and grid-based analyses has numerous computational advantages. However, at least for standard cluster-based methods [e.g., 16], the grid-cells (pixels) themselves become the information-bearing unit. Thus, patterns that unite species based on similarities in their total ranges may actually become harder to detect [20]. For example, species that shared a historical “center of dispersal”, but dispersed out of that area to different extents might show very different overall ranges around a common area [e.g., 19], and the extreme cases of large and small ranges within such a group of species will only show congruence with species with intermediate ranges [22]. Thus, the detection of these range gradations can only be made through the extension of comparisons beyond direct congruences, through higher-order spatial relationships in network approaches [e.g., 23], or through indirect spatial congruences, as proposed here (see Methods).

Methods that equate the frequency of co-occurrence with its relevance will fail to detect unusual, seemingly quirky cases of shared ranges by only a few species that may, nevertheless, represent important or interesting natural phenomena. An intermediate approach to recover such small units is presented by the Endemicity Analysis [24,25]. It searches for sets of grid cells matching the distribution of as many congruent species as possible (in practice, the same as testing congruences to hypothetical species distributions) [26]. Much like other grid-based systems [e.g., 15,16,25] it is subject to scale bias (e.g., pattern detection varies according to grid-cell size) [26,27].

Another difficulty in biogeographic pattern recognition is that an excess of information can create “noise” in analyses and actually hide relevant patterns. Much as uninformative genetic loci must be culled from phylogenetic analyses of relatedness, the presence of widespread species, with little or no biogeographic information to offer, may distort biogeographic relationships [29]. Thus, filtering out biogeographically uninformative species should improve the performance of any method.

In this paper, we describe Spatial Congruence Analysis (SCAN), a framework that explicitly defines and quantifies spatial congruence between species, and recognizes groups of congruent ranges as relevant biogeographical units, representative of potentially interesting natural processes. By allowing different degrees of congruence in a network of species spatial relationships, this approach permits indirect comparisons and so should recognize gradients of species ranges. By using entire ranges as the unit of comparison, rather than grid cells, problems of range distortion are minimized and focus is maintained on relationships among species’ ranges rather than arbitrary spatial units. Finally, by detecting natural congruence-based groups, biogeographically uninformative species can be identified, as can independent overlapping patterns, based even on very few species. The method is first presented with a hypothetical dataset to exemplify its behavior and specific metrics. Pros and cons of the approach are then further illustrated with real bird species ranges and compared to previously published analyses or results using other available methods.

## Material and methods

### SCAN overview

SCAN is based on a script written in R environment [30], which relied mostly on *sf* package [31] for spatial tasks (**S2 Script**). In a nutshell, it uses a quantitative measure of congruence (see below) to compare polygons of species ranges. Comparison of a chosen initial species with all others yields a set of species whose ranges are spatially connected at a given threshold of congruence. Subsequently, each of those already grouped species is also compared to all others, sometimes accruing more species to the list. The process is repeated until no more ranges are found to be congruent at the threshold being evaluated. This group is considered “closed” and is referred to as a “biogeographic element”.

The process begins at the highest congruence threshold (e.g. 100%), saves the results of the iterative comparisons (if any congruence is found and if the group closes), and then reinitiates using the same initial species for the next round with a lower threshold (e.g. 99%). Each time a new biogeographic element is detected (i.e., that level of congruence leads to closure), this new list of species is saved. This process continues anew at each progressively lower congruence threshold until the comparison no longer leads to closure. The set of all biogeographic elements (closed lists) at all congruence levels is considered the “biogeographic complex” (see below) of that particular reference species.

### Congruence index

Determining spatial congruence is not a trivial problem because ranges may differ in position, area, and shape. Two ranges of equal area and shape may vary in the amount of overlap, just as two ranges of equal shape and central position may differ in size, and yet areas of the same size and position may overlap only slightly if their shapes are very different (Fig 1A). To distill these differences into a single index, spatial congruence between two species can be calculated by the product of area of overlap weighted by the relative area of each (Eq 1). This generic spatial index was proposed in the “Goodness of Fit” method to compare maps of Hargrove et al. [32] and is hereafter referred to as the “Spatial Congruence Index” – C_S_, defined as follows:

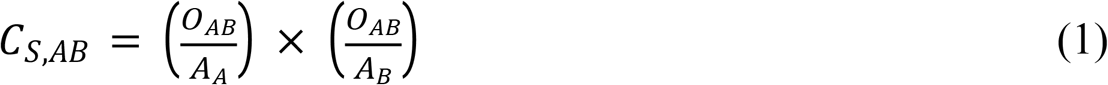

where C_S_ is the spatial congruence index, O_*AB*_ is the area of overlap, and A_*A*_, A_*B*_ are the areas of the ranges of species *A* and *B*, respectively [32]. The index varies from zero (no congruence) to one (perfect congruence) and can also be expressed as a percentage. It allows the use of vector-based distributions (polygons), thus avoiding the scale-dependent shape distortion caused by the use of coarse grid-cells [33]; however, this index can also be used with grids. Furthermore, any other quantitative congruence index may be substituted in the method here proposed, if preferred or if required to compare other range representations, such as the probabilistic surfaces resulting from species distribution models.

**Figure 1.**
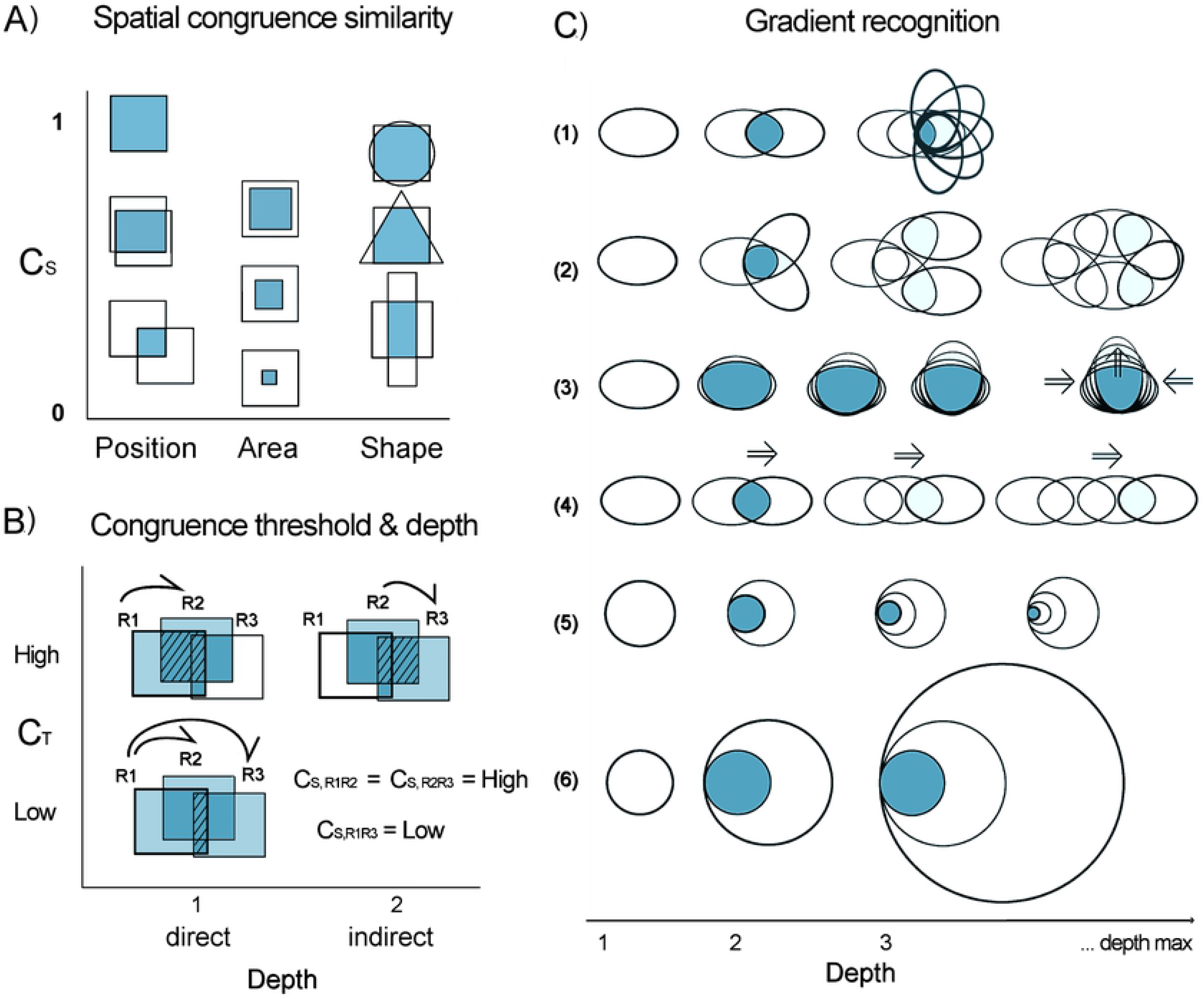
Schematic representation of the theoretical behavior of spatial congruence with respect to range similarity, threshold values, depth of direct and indirect congruences, and gradient recognition. All figures are spatial representations; darker tones indicate shared ranges. (A) The C_S_ index (y-axis) compares position, extent, and shape between two species ranges. For identical size and shape, position determines the amount of overlap; when shape and position (e.g., centroids) are the same, one area will reside entirely in the other and difference in area alone will determine their congruence; and, finally, if area and position are the same, shape will be determinant. (B) The congruence threshold (C_T_) is the reference for spatial relationships. Range R1 is directly congruent with R2 when their C_S_ (calculated from their area of overlap – hashed area) is greater than or equal to C_T_. Indirect relationships occur when ranges are liked by congruence in a concatenated chain of C_S_ ≥ C_T_. R1 is indirectly congruent with R3 because R2 is as congruent (same C_S_) to R3 as it is to R1. At relaxed threshold requirements (i.e., C_T_ < C_S,R1R3_), R2 and R3 are both directly related to R1. (C) Indirect links allow the recognition of many types of syndromes of shared distributions potentially associated with distinct historical or ecological range drivers. Links may, at least theoretically, lead to nuclear regions of congruence (example1), congruence zones following patches of favorable habitat (e.g. mountain slopes, or riverine habitats; ex. 2), gradients of expansion or contraction (ex. 3), or linear gradients (ex. 4). Indirect congruence (depth) at very low congruence thresholds may also lead to scalar distortions (ex. 5,6).

### Direct versus indirect congruences

Spatial congruences, here, are always evaluated with respect to a specific congruence threshold (C_T_). Two species are directly congruent when their calculated C_S_ is greater than or equal to a particular C_T_ (Fig 1B). Indirectly related species are those linked through a chain of direct connections using a third linking species. For example, if species ranges *A* and *B* are directly congruent at a given C_T_, and *B* and *C* are also directly congruent, then *A* and *C* are indirectly congruent at this C_T_, using *B* as a link (*A↔B↔C*). These direct and indirect relationships are recovered and organized by the algorithm at every referential C_T_ value. Thus, the method begins setting the current C_T_ and comparing the spatial distribution of a reference species to the range of all other species. At each C_T_ analyzed, any species with C_S_ ≥ C_T_ is directly congruent. The next pass involves comparing each of these directly congruent species to all other species. Any additional species added in this second pass (and all subsequent passes) are indirectly congruent to the reference species at the same C_T_. The number of passes required such that the species group closes (i.e., no additional congruent species are included) is equal to the number of links to the longest indirect chain and is called the “depth” of that species group (Fig 1B-C). Depth is a metric that emerges from the analyses; however its exact biological significance and its computational utility are not clear, but may be subject to future study.

Including indirect congruences in the recognition of these biogeographic elements has two distinct advantages. First, it allows the recognition of syndromes of shared distributions (Fig 1C). The set of species in a closed group, at any given C_T_, will be very similar to that formed by starting the analysis with other members of the group. Thus, this is a “natural” grouping. Second, indirect congruences allow the recognition of gradual relationships among species ranges (Fig 1C). Although it is conceivable that virtually all species be related to one other through such gradual indirect congruence, this is controlled by congruence threshold requirements. In practice, groups of species sharing biogeographical properties may close even at fairly low C_T_ (see Results).

### Control parameters

All reference species have their C_S_ calculated with respect to all species available in the whole pool of taxa. A list of species to be analyzed as references and maps for all taxa are the basic inputs required to run the algorithm (a tutorial is available at **S1 Script**). The most important analytical settings are: 1) the maximum depth (number of links in the chain of indirect comparisons allowed before stopping the analysis with a given reference species); 2) minimum C_T_ (the lowest congruence threshold analyzed); 3) interval size for relaxation of C_T_ for each subsequent analysis (the default is 0.01). In addition to these three numerical values, the user may stipulate a criterion of spatial overlap. If adopted (default), this criterion requires that a congruent range must include at least some area of overlap with all previously grouped species.

Once these parameters are chosen, the iterative procedure of evaluation of spatial relationships across congruence thresholds will be executed for all references, one by one. At any time, the analysis will stop at a given C_T_ with a given reference species when a group closes, that is, the next depth of indirect comparisons adds no new species. That closed group is recorded and the comparisons begin again at the next lower C_T_. The entire sequence of comparisons for a given reference species stops when groups no longer close before reaching the stipulated maximum depth, or the minimum congruence threshold, or do not meet the spatial overlap criterion. When the stopping point is reached for a given reference species, another species is run to completion, and so on until all reference species have been analyzed.

### Biogeographic elements and complexes, synonyms, and metrics

A closed group of species whose ranges are congruent at a particular C_T_, here called a “biogeographic element”, is analogous to Hausdorf’s “biotic element”: that is, “a group of taxa whose ranges are significantly more similar to each other than to those of taxa of other such groups” [22]. Here, any biogeographic element will always be dependent on a specific, explicit congruence threshold and reference species, and will also be associated with a particular depth (see **S1 Fig**). The biogeographic element, thus, is a suite of species. The spatial area determined by their distributional congruence, equivalent to chorotype [4], can be delimited in different ways. This method reveals both a *“*total area”, which congregates the ranges of all grouped species, and a “common area”, defined by their spatial intersection. One simple metric, the “ratio between common and total areas” (intersection/union), ranges from 0 to 1 and addresses the trade-off between spatial comprehensiveness and cohesion (see Discussion).

SCAN differentiates those reference species forming biogeographic elements, “informative species”, from those not leading to closed groups, at any congruence threshold, “non-informative species”. For any informative reference species, the set of all biogeographic elements over the range of congruence thresholds constitutes a “biogeographic complex”. “Synonymous” elements or complexes (or simply “synonyms”) are those derived from distinct reference species that converge on the same group of taxa. Nested patterns are those composed by subsets of larger patterns. Parameters such as maximum and minimum C_T_, depth, and derived metrics, such as the number of species, can be used to identify synonyms and compare elements or complexes derived from distinct species or regions (**S1 and S2 Tables**). Although synonyms and nested complexes are overlapped by definition, partial spatial overlaps may also occur between independent elements, with no species in common (see Results). Synonymous complexes necessarily are generated by different reference taxa; here, they are represented by and named after the reference taxon with the highest mean congruence (C_S_) with all other species in the complex (**S3 Table**).

### Applications of the method

To explore and illustrate this protocol we use two datasets, one hypothetical and the other of real bird species distributions. First, a hypothetical set of species ranges was used to evaluate the capacity of SCAN to detect gradients. In addition, the effects of maximum depth settings over pattern detection were explored (see S1 Script). The problem, proposed by Kreft and Jetz [34], is when two seemingly distinct sets of species (for example a northern and a southern groups) contain a succession of species with ranges each slightly greater than the previous such that those with the most extensive ranges in each group actually overlap partially the widest-ranging species of the other group (**Fig 2**). Any biogeographical method would be seen to fail if it did not recover the two groups as separate congruent units or, worse, if it identified the transition zone between them as an independent biogeographical unit, which is, obviously, a false conclusion from a biogeographical perspective.

**Fig 2.**
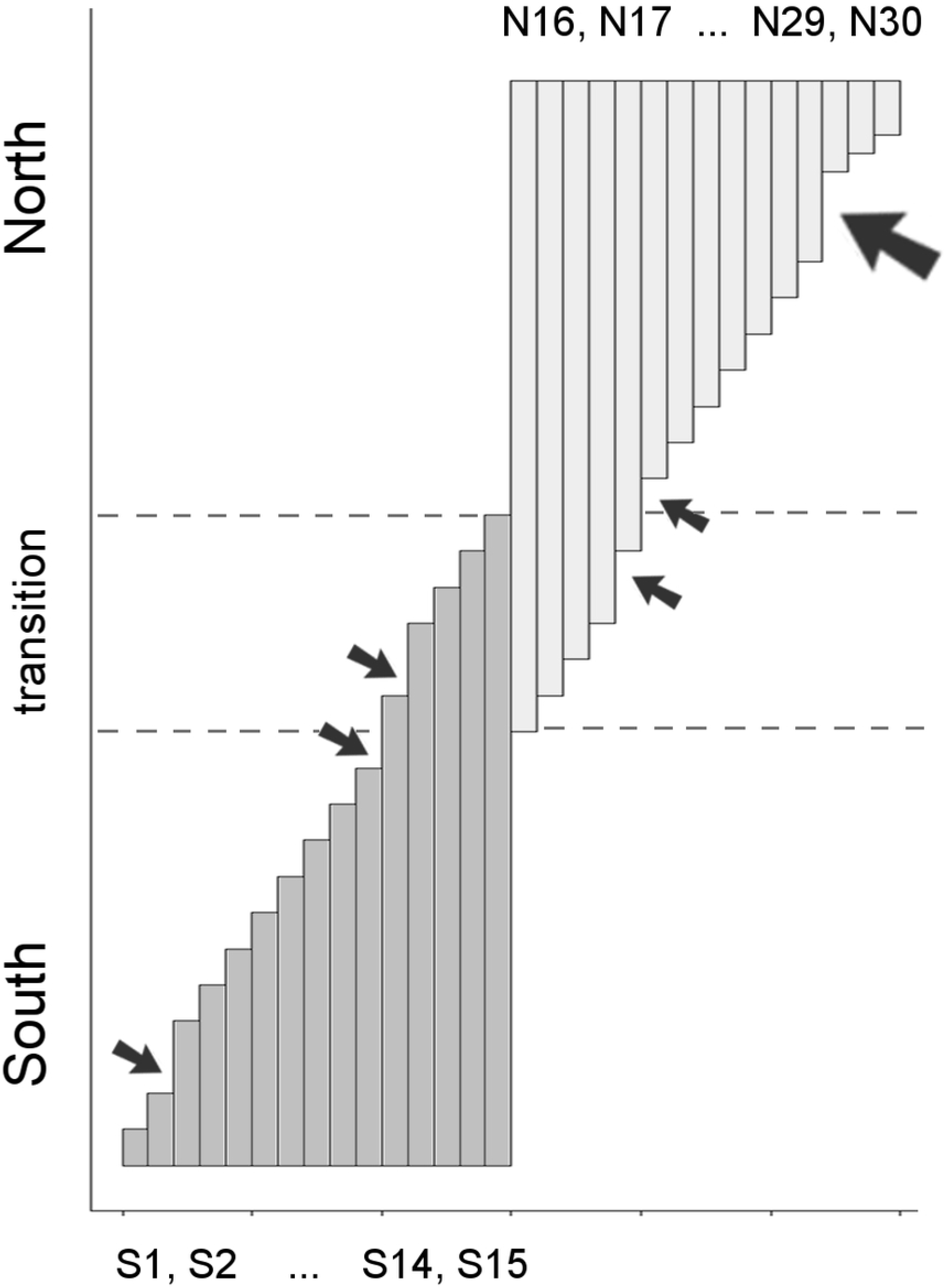
Schematic representation of the hypothetical gradient of thirty species ranges proposed by Kreft & Jetz [34]. Two well delimited biogeographical centers of diversity, a northern (pale gray) and a southern (medium gray), have species extending their ranges through a zone of transition, which is recognized erroneously by some methods as a distinct center. Each vertical bar represents the latitudinal extent of the range of a particular species (named S1-15 and N16-30). The longitudinal ranges (bar widths) of all species are assumed to be identical, such that they all overlap spatially. The original symmetric scheme [34] included a few small exceptions (small arrows) to the otherwise uniformly graded range differences. To further examine the influence of non-uniform gradations on detection of congruence patterns, we reduced the range differences among N28-30 relative to the others, which adds a single slightly larger range discontinuity (large arrow).

For subsequent exploration of the method, a dataset with real species distributions from BirdLife & HBW [35] was used to examine the influence of reference species on resulting biogeographic elements, to examine the relationships between parameters such as C_S_, C_T_, and depth, and to illustrate the method with maps of real patterns detected. The set of taxa analyzed as reference species is here defined arbitrarily as those 1095 birds for which at least 50% of their distribution is within Amazonia [sensu 33]. To allow recognition of possible patterns that extrapolate the study area (Amazonia), those species’ ranges were compared to a larger dataset containing all 3083 New World bird species whose ranges overlap the Amazon even only slightly.

As mentioned before, in addition to detecting patterns of congruence among species, SCAN also identifies those reference species that show no meaningful spatial relationships with others, at least according to the criteria proposed here. Unless further taxonomic or distributional updates become available, these species ranges may be thought of as biogeographically uninformative. To test the effect of inclusion of uninformative species on methods of spatial classification that do not make this distinction, we ran analyses in Infomap Bioregions (hereafter Infomap; https://www.mapequation.org/bioregions), a modern framework incorporating recent developments of network theory to bioregionalization [23,37]. The Infomap analysis was conducted on three versions of the Amazon bird dataset: all 1095 species, the 610 Amazonian “endemics” (at least 90% of their range within Amazonia), and those 329 species identified as biogeographically informative by SCAN. This last set included only those species that formed biogeographic elements within Amazonia, i.e. not largely including other biomes (e.g., Atlantic Forest, trans-Andean regions) or encompassing the entire Amazon. To take advantage of the adaptive spatial resolution in Infomap [37], we adapted settings to Amazonian standards of species richness to maximize pattern recognition (‘max_cell_size’ = 1, ‘min_cell_size’ = 0.5, ‘max_cell_capacity’ = 200, ‘min_cell_capacity’ = 4), and default values (4, 1, 100, 10, respectively). Infomap’s ‘cluster costs’ used were 1 (default), and the more relaxed 0.97 and 0.94 settings.

## Results

### Gradient simulation

Analysis of the simulated gradient resulted in a total of 23 unique biogeographical patterns (**S1 and S2 Tables; Fig 3**). SCAN always correctly recognized north and south as distinct groups. The number of informative species varied according to max-depth settings (**S2 Table**). At the shallowest depth limit (maximum depth=3), approximately half of the community gave rise to biogeographic elements. At the extremely relaxed max-depth=10, all species expanded their indirect congruences to encompass either the entire southern or northern groups, but never included species from the opposite group if the “spatial overlap criterion” (see Methods) was implemented. Intermediate scenarios with alternative and plausible classification schemes were achieved at depth limits of 5 and 7, but always nested within the overall north-south distinction (**Fig 3**). These alternative configurations are ideal to show how the C_T_ relaxation allows the incorporation of slightly less congruent species, at each lower C_T_ round (direct) or extended depth (indirect).

**Fig 3.**
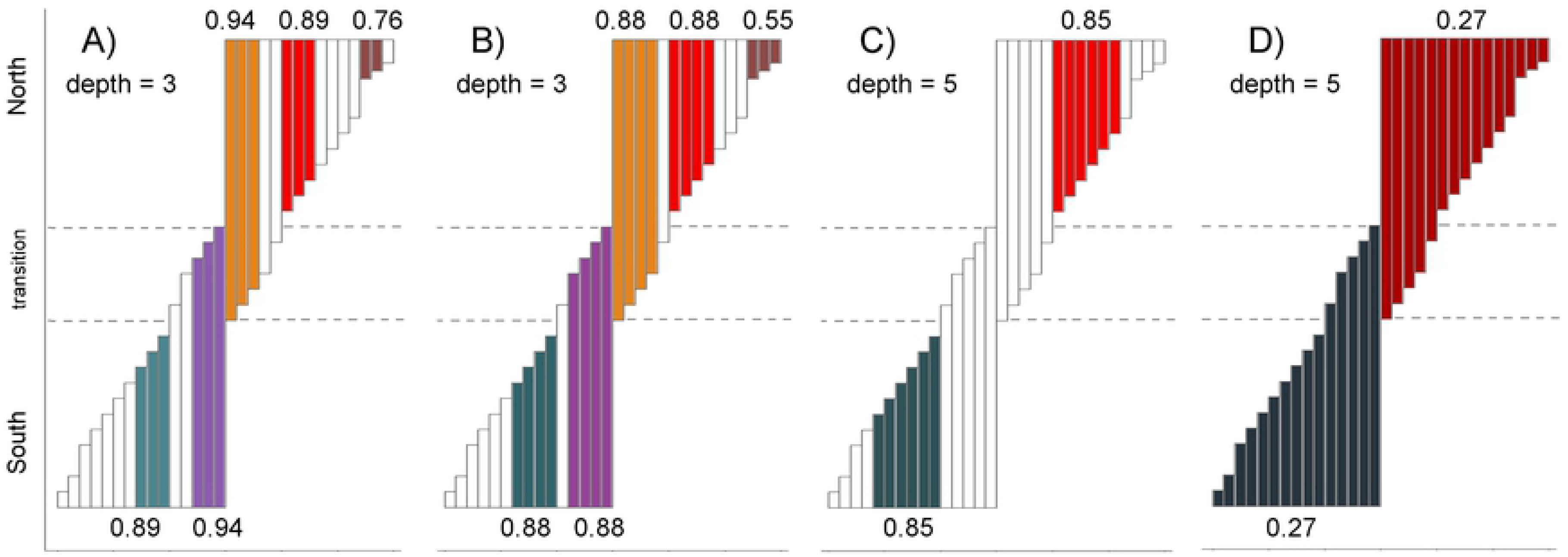
Biogeographic elements detected in a simulated gradient of ranges. Like colors indicate species ranges (bars) grouped into biogeographic elements recovered at particular congruence thresholds (minimum C_T_ shown at the base of each element). White bars indicate “biogeographically uninformative” species that did not give rise to and were not included in any biogeographic element, at each corresponding maximum depth setting. Four examples are shown (see **S1 Table** for full details). A) Small but highly congruent patterns were located next to zones of range discontinuity and favored larger ranges, with smaller proportional area differences. The extended discontinuity at the northern extreme allowed the recognition of one element composed of small-range species. B) At slightly lower congruence thresholds, elements included adjacent species with smaller ranges. C) Increasing depth (indirect congruence) increased element size at relatively high congruence thresholds. D) At lower congruence thresholds, the entire northern and southern elements were recovered. Allowing for extensive indirect range comparisons (maximum depth of 10, not shown), most reference species led to convergence on the whole community. Note that all recovered elements can be nested in others and there was no non-nested overlap among elements nor recovery of the transition zone as a biogeographic element.

### Bird dataset

Applying SCAN to (or “SCANning”) the entire 1095-species Amazonian bird dataset resulted in a very large suite of biogeographic elements and complexes with interesting implications for the study of Amazonian biogeography. That subject goes well beyond the objectives of this paper and will be presented in detail separately. We examined the overall performance of the method running exploratory analyses in R to synthesize relationships between parameters and metrics. These analyses used settings preliminarily defined after pilot tests, which showed stable and consistent results: 0.01 as the interval between C_T_ rounds, 0.1 as the min-C_T_, and 7 as the max-depth for the bird dataset (although this limit is rarely reached; see below). Specifically, the cases of *Icterus nigrogularis* (Yellow Oriole) and *Amazilia tobaci* (Copper-rumped Hummingbird), at the northern extreme of the Amazon (**Fig 4**), and a genus of hummingbirds (the “brilliants”, *Heliodoxa* spp.), with a large variety of habitat associations and peculiar ranges (**Fig 5**), are sufficient to demonstrate here the main properties and characteristics of SCAN for real life distributions.

**Fig 4.**
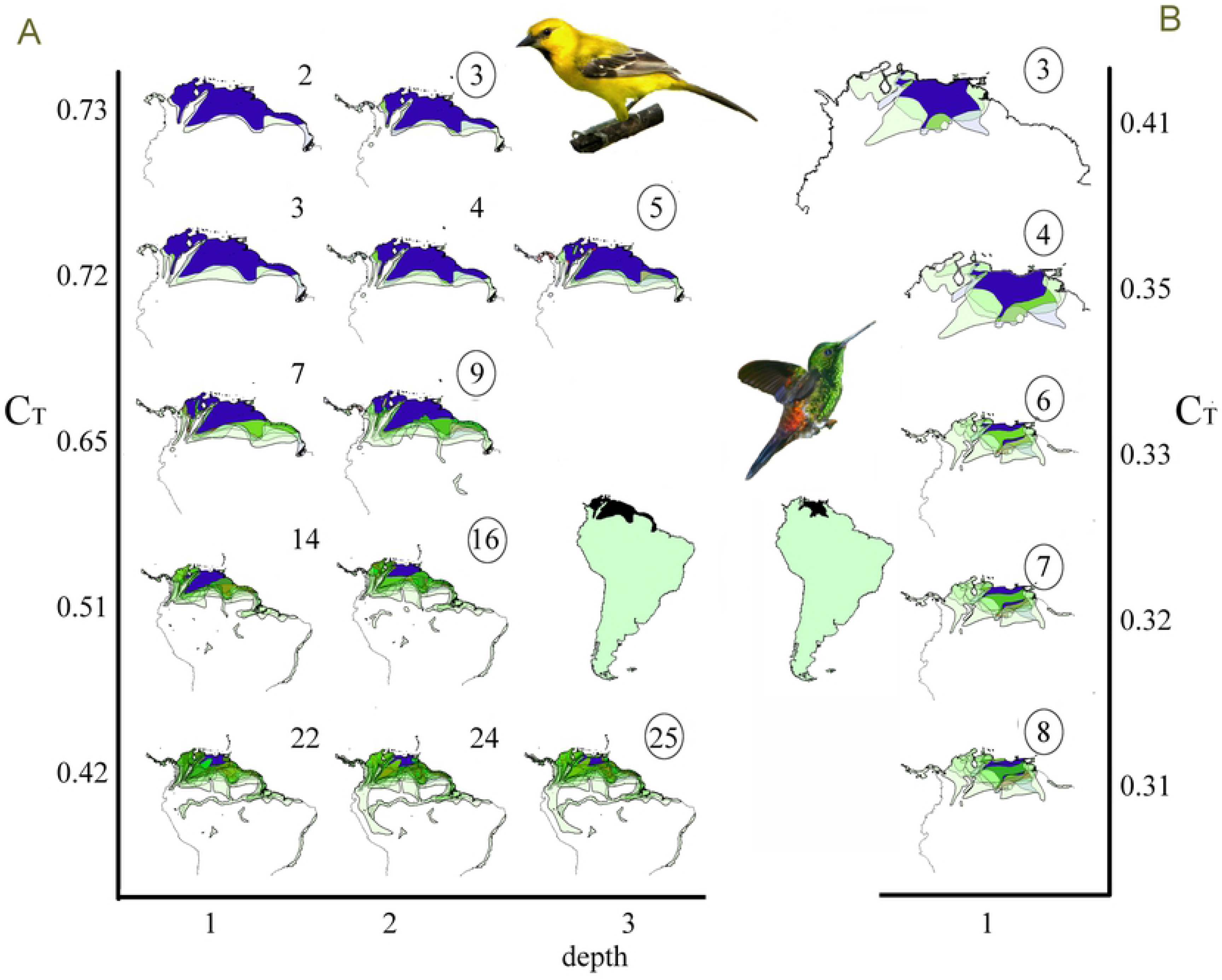
Illustrative whole biogeographic complexes of *Icterus nigrogularis* and *Amazilia tobaci* bird species. Diagrams depicting congruence threshold (C_T_, y-axis, free scale) and depth (x-axis) relationships are presented. (A) *Icterus nigrogularis* (Yellow Oriole) and (B) *Amazilia tobaci* (Copper-rumped hummingbird). For each C_T_ and depth, ranges (light-green) are overlaid; number of species in each group are shown for each round at each depth. Circles depict the closed groups (biogeographic elements) at the end of each round of threshold analysis. The common area is shown in dark-blue. South America insets show reference species ranges as black. Grouped species and their synonyms are listed at S3 and S4 Tables, respectively. Geographic scales are variable as maps are zoomed out to accommodate successively larger elements. Credit: images of *I. nigrogularis* and *A. tobaci* adapted from Wikimedia Commons under Attribution-Share Alike 2.0 Generic license (creativecommons.org/licenses/by/2.0/deed.en).

**Fig 5.**
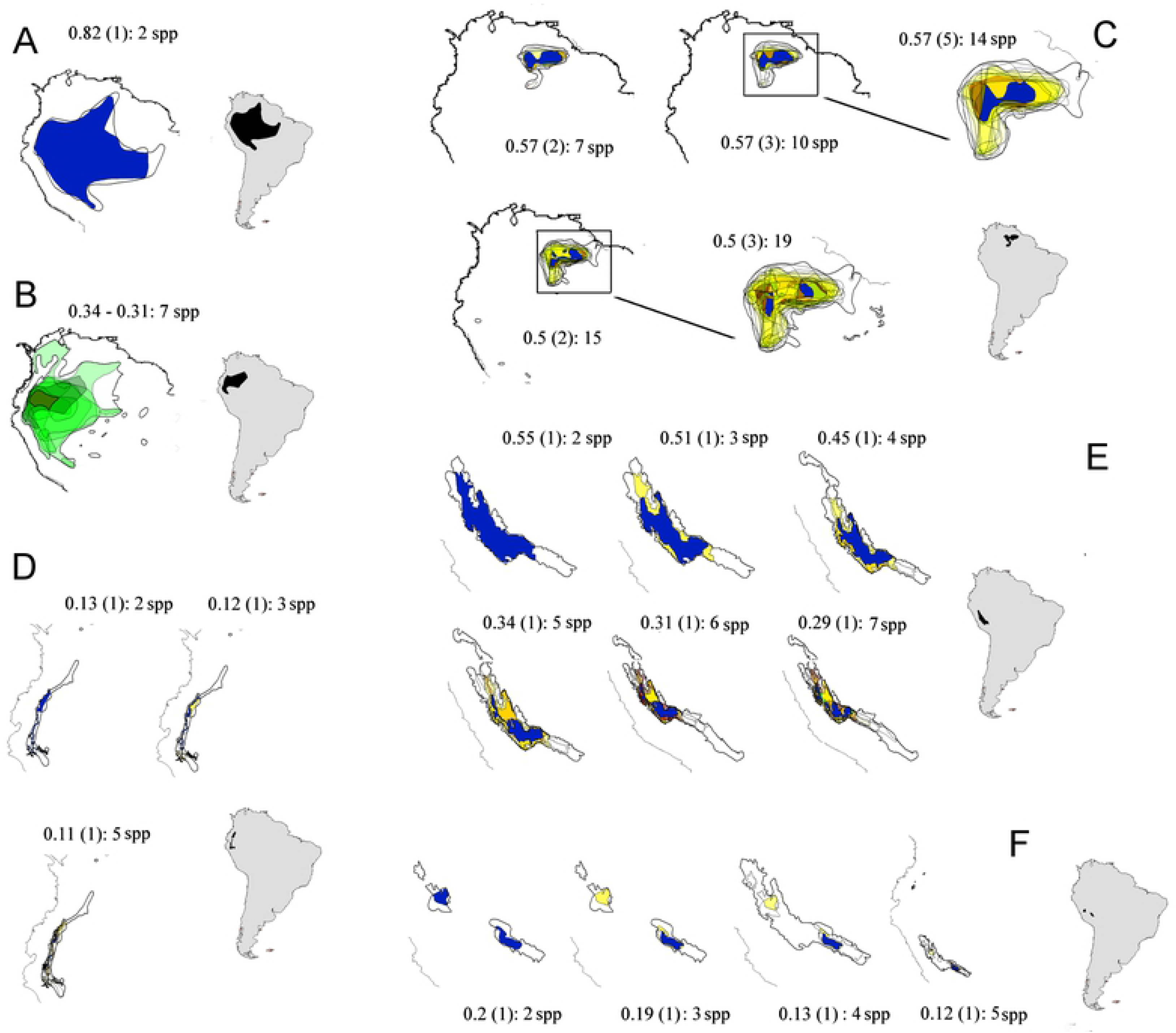
Biogeographic elements of *Heliodoxa* hummingbirds in lowland Amazonia and Pre-Andes. Brilliant hummingbirds include a large diversity of species with distinctive biogeographical patterns. Widespread lowland species (A) *H. aurescens* (Gould’s Jewelfront) and (B) *H. Schreibersii* (Black-throated Brilliant) do not compose biogeographic elements. Highland species show spatially localized patterns; in the east in the Pantepui (C) *H. Xanthogonys* (Velvet-browed Brilliant) and west in the Pre-Andean region: (D) *H. gularis* (Pink-throated Brilliant); (E) *H. branickii* (Rufous-webbed Brilliant); (F) *H. whitelyana* (Black-breasted Brilliant). These summarized views of complexes present the range of congruence thresholds, depth (in parentheses), and number of species as descriptive parameters. Grouped species are listed in **S3 Table**. Ranges of individual species leading to biogeographic elements are shown in yellow; the common area for all species is in blue.

Roughly half of the bird species analyzed as references (558) were recognized as informative by the algorithm. These species gave rise to approximately 150 patterns, including synonyms and nested complexes, grouping anywhere from two to 99 species in an element. Preliminary explorations showed that the algorithm chooses a very limited set of highly congruent range overlaps (**S2 Fig**). Reference species with larger areas had higher C_S_ means (i.e. the average C_S_ between one species and all its overlapping ranges). Range areas and C_S_ means had no effect on the number of species (richness) in patterns detected, but relaxed congruences and depths led to patterns with greater species richness (**S3 Fig**). The full set of graphic preliminary explorations is presented as Supplementary Material (**S2 and S3 Figs)**.

Biogeographical complexes derived from this real-life bird community were highly diverse in many aspects (**S1 Table; Figs 4 and 5**). The simplest patterns had only one element with a fixed group of species, usually in a very limited range of threshold values, as shown by the lowland hummingbird *Heliodoxa aurescens* (**Fig 5A**). Alternatively, “shallow” patterns added species across a range of congruence thresholds but showed no indirect connections, grouping species only at the first depth level (**Figs 4B and 5D-F**). The depth ‘dimension’, as predicted, revealed gradients through indirectly related ranges. This potential was elegantly illustrated by the *Icterus nigrogulari*s complex, which gradually, range by range, extended its total area through the Guianan coast and subsequently reached the central Amazon via the Amazon river channel (**Fig 4A**).

SCAN can recognize more than one biogeographic element centered on essentially the same general area, but with considerably different spatial limits and containing mutually exclusive sets of species, as demonstrated by *I. nigrogularis* and *A. tobaci* (**Fig 4**). Similarly, the *H. xanthogonys* pattern grouped highland birds of the Tepuis region that are completely overlapped (at this geographic macro-scale) with birds typically associated with lowland patterns (**Fig 5C**). Other *Heliodoxa* hummingbirds illustrate SCAN’s accuracy recognizing patterns derived from species with small, somewhat linear ranges, sometimes at very low congruence thresholds based on subtle similarities of range position and shape, even grouping species with similar disjunct distributions (**Figs 5D-F**).

### Non-informative species and spatial classification

Bioregionalization in Infomap led to very different results depending on how many species were included. For the first set (all 1095 species), even with the astonishingly high number of 440 bioregions identified at a relaxed cluster cost (0.97), no internal divisions in Amazonia were recognized (**Fig 6A**; see details in **S4 Fig)**. Instead, the bioregions recovered were mainly associated with range limits of widespread species, far from Amazonian borders (**Fig 6A**). With the ‘endemic’ set (609 species) Infomap recovered internal divisions within the Amazon. Many units, however, were mostly overlapped patterns with few variations in extent, punctuated by scattered idiosyncratic bioregions, particularly in areas of ecological tension (e.g., Tepuis). Infomap also emphasized patterns at the limits of widespread species (See Chocó and Atlantic Forest regions; **Fig 6B**). When given the optimized group of 329 limited-range lowland species identified as biogeographically informative by SCAN the results stabilized across distinct scale and cluster cost settings, and showed a well known pattern based on species turnover across major Amazonian rivers [10,38]; **Fig 6C** and **S4 Fig G-I**).

**Fig 6.**
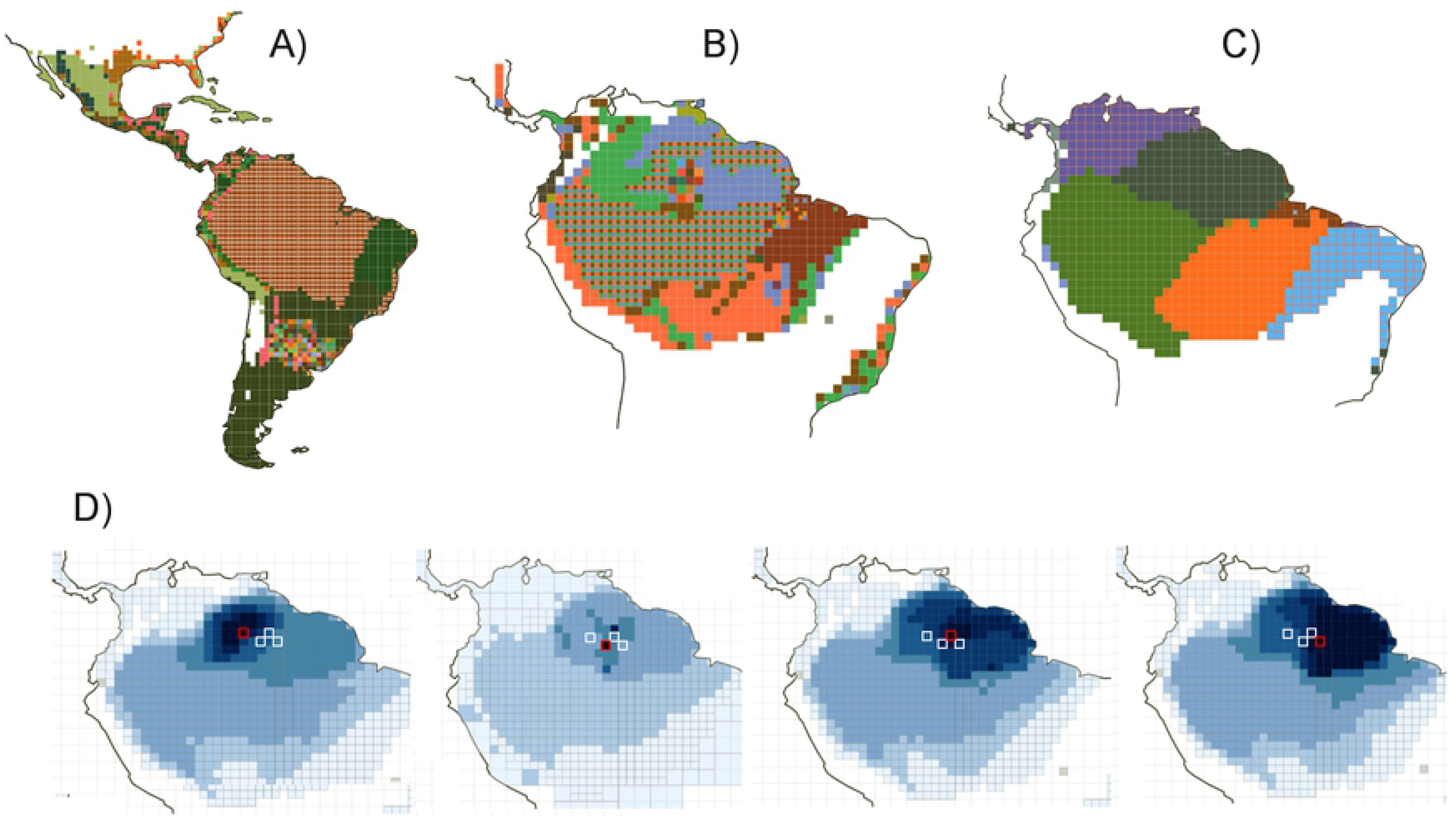
Too many species for bioregionalization? (A) Infomap Bioregions [37] cannot recover internal biogeographical structures in the Amazonian lowlands for large datasets, irrespective of scale and cluster settings (see more scale and cluster options in **S4 Fig**). (B) Even using only species with more than 90% of their distributions restricted to Amazonia there are redundant widespread bioregions and variable pattern configurations across scale and cluster settings. (C) Results based on a subset of biogeographically informative species with ranges relatively restricted within Amazonia, as identified by SCAN. This subset revealed a stable configuration across scale and cluster settings. This classification matches previous species turnover assessments [38]. D) Infomap uses patterns based on networks of species inter-relationships centered in grid-cells which are analogous to the biogeographic elements described here. The draw-back of this cell-based approach is that each cell has its own idiosyncratic pattern. Even contiguous cells may show very distinct representative species (see **S5 Fig** for details), and often mix species belonging to distinct ecological and biogeographical domains as indicative species (e.g., Tepuis, lowland *terra firme* forest, *várzea* floodplain specialists)[39,40].

The characteristic relational web of ranges based on grid-cells presented by Infomap resembles, and is probably analogous to, the biogeographic elements identified by SCAN (**Fig 6E**). Specialized birds clearly belonging to distinct biogeographic niches, such as highland, riverine specialists, or lowland *terra firme* birds, are mixed as indicative species in these cells, while adjacent ones often give very distinct results (**Fig 6D** and **S5 Fig**). Unfortunately, these cell-based patterns are totally idiosyncratic, i.e., each cell has its unique pattern, which hampers the direct interpretation of these relationships in biological or geographical terms (**Fig 6E** and **S5 Fig**).

## Discussion

SCAN proposes a comprehensive description of biogeographical scenarios based on the whole web of spatial interrelationships of species in a community. As a method inspired by biogeographical concepts, it has many unique properties, combined for the first time. Explicit pairwise congruences permeate all steps of the analysis. To Linder [22], bioregionalization approaches should be oriented mainly by spatial congruence, although he recognized the difficulty associated with this objective. SCAN then assesses the informative potential for each species separately and so allows the investigator to decide whether their inclusion in subsequent analyses is justified. Also, the algorithm does not classify grid-cells based on their similarities with other cells (e.g., cluster, ordination), but recognizes mutually congruent groups (biogeographic elements) in a network of species linked (and weighted) by spatial congruences. Degree of congruence (congruence threshold) is an intuitively simple concept likely to have biological relevance. A community under more spatially precise, long-lasting, or intense spatial drivers should have more congruent ranges. Nevertheless, there is no theoretical basis for establishing any specific numerical threshold, and the method permits exploring alternatives. Biogeographic elements are groups of species, not specific regions, and the delimitation of the geographic area they represent can be determined by the investigator based on the specific questions under study. Furthermore, all these characteristics allow the identification of patterns overlapped in space but with distinct species compositions. Although some authors argue that areas of endemism cannot co-exist in space [e.g., 39], there are compelling theoretical [22] and practical [27] arguments for recognizing spatially coexisting biogeographic patterns. Finally, SCAN correctly recognizes and describes gradients of range distributions. Despite the key role gradients have had in the development of ecological thinking [42,43], they have not received the same attention in the field of biogeography [but see 15,42,43], which tends to see imperfect congruence as an inconvenient deviation from idealized responses to geographic barriers.

The main characteristics of the method were demonstrated by the data sets analyzed. Most importantly, these analyses showed that SCAN generates results that may be indicative of possible spatial drivers of species ranges, and can be used to make biogeographical decisions. The richness in details and alternative results of the simulated gradient reinforce the idea of a high “biogeographical resolution” of SCAN results. The bird dataset showed how patterns derived from real life species are idiosyncratic, presenting highly distinctive alternative configurations. This apparent contradictory interpretation is better seen as evidence for the co-existence of distinct but complementary phenomena driving spatially related species. Highly variable complexes, for instance, congregating from small highly congruent elements to large spatial gradients recovered under relaxed thresholds, may carry information about both vicariant causes, and processes responsible for pattern deconstruction, such as local extinctions, dispersal events, and differential responses controlled by species-specific traits [46,47].

Many of SCAN’s properties are derived from the objective focus on species ranges and their spatial congruences; these relationships are crystallized in SCAN’s one-layered networks, where species are vertices connected by their respective spatial congruences (edges). The algorithm proposed here to traverse the network is a type of “Breadth-First Search (BFS)” self-stabilizing spanning tree [48], in which all vertices connected to a reference (species) are analyzed at the first depth level before starting the analyses of indirect connections, at each subsequent depth level. This level-by-level approach characterizes BFS, as opposed to “Depth-First Search”, which explores branch connections in full depth before analyzing all neighbors in the same level. In SCAN edges (connections) are weighted by C_S_ estimates. The algorithm, however, evaluates only a subset of the whole network (not the full set of connecting overlaps as in other methods) composed of pairs of ranges matching the congruence threshold being evaluated. In practice, each round analyzes only the direct and indirect links which are “activated” at this respective C_T_. This conditional connection assures that spatial congruences based on whole ranges are considered during algorithm processing, and that all potentially connected species have comparable levels of spatial congruence. When a group closes, the resulting elements are biogeographical units built over pure species-spatial inter-relationships, analogous to the concepts of ‘community’ or ‘module’ in network analysis, that is, sets of nodes that are more densely connected to each other than to other nodes of the network (Rosvall & Bergstrom, 2008)[49]. Each biogeographic element, as presented here, matches the definition of biotic element given by Hausdorf [22]. However, SCAN expands the concept and shows that diverse alternative configurations obtained at distinct thresholds also match the same definition.

### Patterns, parameters, and biological correlates

Examining the biological meaning of SCAN’s parameters calls attention to important trade-offs in pattern recognition. For example, high species richness in a spatial pattern, that is, its inclusion of numerous species, is a desirable feature for any pattern. However, it is usually achieved at the expense of spatial coherence, because more species are usually grouped with lower congruences or greater depths, which in turn lead to smaller common and larger total areas. In communities dominated by idiosyncratic ranges, small gains in species may quickly disfigure the spatial coherence of a pattern. Conversely, a community shaped by effective barriers (Hazzi et al., 2018)[50] or by zones of intense environmental filtering (Blonder et al., 2015)[51], should display patterns that add species while maintaining a certain level of spatial coherence. Highly congruent biogeographic elements can be directly compared to areas of endemism (Szumik et al., 2012)[52], or bioregions (Vilhena & Antonelli, 2015)[13].

The concept of depth appears to have no clear biological analog. In practice, however, it allows the identification of gradients through indirect spatial relationships, in a delicate equilibrium. As seen in the simulated gradient, the chain of links must be limited by a discontinuous interval, which closes a group. This paradox (connection vs. disconnection) is the essence of SCAN: regardless of the congruence threshold, if there are closed groups, then there is spatial cohesion among their constituent taxa relative to the pool. Under low congruence thresholds and large maximum depth settings, however, links may potentially lead to distortions in range sizes, which often prevent a group from closing.

### Concluding remarks

The development of this method was stimulated in part by the desire to minimize assumptions, particularly those related to the biological causes of spatial congruence in ranges and of pattern formation (e.g., vicariance, dispersal, ecological determinism). However, like any model that sets out to describe real-world patterns, some general assumptions apply to this framework. We recognize three. First, species ranges and their direct and indirect relationships are biogeographically meaningful; this is an assumption of all biogeographical analyses. Second, range maps are good representations of such distributions, at least at the spatial scale of interest; this is a potential weak point, in that current range maps in certain regions are notoriously flawed [53]. However, poor distributional data, especially data suffering from chronic omission [e.g., 52], will also affect all other methods, and improvements in maps and taxonomy can be expected over time. Third, the set of analyzed species is a representative sample of the community under evaluation. Adding or removing species will readily influence detection of connections or breaks in congruence through indirect links. Thus, it is important that the species pool analyzed be made explicit.

Perhaps the most important take-home message of this paper is a cautionary ‘step-back’ regarding the generic use of bioregionalization methods for spatial classification, recognition of biogeographical patterns, and assessment of their historical and ecological drivers. In some cases, spatialized biological information may be being discarded. Conversely, the excess of idiosyncratic range information may bias classification schemes of standard methods. Common biases and assumptions may even obscure the effects of distinct but complementary drivers of patterns of species distribution.

The richness of patterns detected by SCAN is a reflection of natural phenomena and natural groupings of species. However, the complex webs of spatial interrelationships revealed require more effort to organize and interpret than results of other methods. The description of biogeographical scenarios depends on posterior identification of synonymous and nested complexes, for example, as encountered in both datasets examined in this study. Rather than a drawback, this post-algorithm analysis can be viewed as an opportunity to better understand these patterns and their variations, species properties, distinct spatial outlines and their environmental correlates, and to make explicit the objectives and assumptions of a particular investigation. For simple complexes the interpretative options are limited (e.g., **Fig 3**). For larger ones, the amount of complementary information may constitute a real challenge (e.g., **Figs 4 and 5**). SCAN asks the biogeographer to interpret individual complexes as a suite of plausible biogeographical scenarios. Highly congruent elements highlight potential barriers or sharp ecological limits. Relaxed patterns may reveal taxonomic or environmental correlates of selective permeability of barriers, and potential dispersal routes. Trade-offs and criteria of spatial coherence and comprehensiveness may also be complemented by biological or environmental information to support both spatial classifications and assessments of spatial drivers. The approach proposed here describes patterns of spatial congruence – the burden of biogeographical interpretation and attribution of processes to explain the patterns is independent of the analysis and lies with the investigator.

## Acknowledgments

This paper is dedicated to Cláudio J Barros de Carvalho. We are grateful to Camila F Ribas for valuable comments on earlier versions of the manuscript, and all the staff at the Programa de Pós-Gradução em Ecologia at Instituto Nacional de Pesquisas da Amazônia - INPA. Joel Cracraft and Thiago SF Silva kindly criticized many of CAFRG’s early unconventional ideas and hosted CAFRG at their laboratories at the American Museum of Natural History - AMNH, NY, and Unesp-Rio Claro, São Paulo, Brazil, respectively. This paper was inspired by the distinct but complementary interpretations of patterns of bird distribution in Amazonia of Jürgen Haffer and Joel Cracraft. We have no conflicts of interest.

## Author contributions

Conceived the method, experiments, and analyzed the datasets: CAFRG. Wrote the paper: CAFRG and MCH.

## Supporting information

**S1 Fig. Schematic overview over the spatial congruence framework.** (A) Spatial representation of a hypothetical community of ten simulated species (*c1-c10*). (B) Congruence similarities between all overlapping species pairs (C_S_ ≥ 0.01) are used for all direct and indirect comparisons in the following steps. (C) At relatively high C_T_, as references species, *c1* and *c6* each gives rise to only one biogeographic element composed of two directly-related species (C_T_ = 0.67 and 0.71, respectively). (D) Depending on C_T_, *c4* may have numerous indirect relationships. At thresholds (C_T_) between 0.46 and 0.22, it is only (directly) connected to *c9*. As of 0.21, *c4* is also directly connected (depth 1) to *c8* which links (indicated with arrow) to *c6* (depth 2), which in turn links to *c7* (depth 3). This biogeographic element (indicated with an asterisk) will be further examined in subsequent examples. At 0.19, an additional link, between *c9* and *c3*, appears at depth 2. However, with this suite of ranges, there is no longer any area of overlap among all ranges (shown in black when it occurs); the range overlap would cause the analysis to drop these round altogether. (E) All informative reference species and their respective biogeographic elements recovered (colored as in preceding figures). Ranges *c5* and *c10* were not included in any elements, and *c3* was included in some groups, but did not give rise to biotic elements. The element [*c4+c9+c8+c6+c7*] can be derived from any of its constituent species (dark blue; asterisk). (F) Possible classification schemes show the trade-off between congruence and comprehensiveness. A highly congruent scheme has three independent two-species elements (blue, green, and red patterns), and *c3* and *c8* are out (yellow). A less congruent scheme (asterisk) has a very comprehensive pattern (blue) based on less congruent indirect links, with a common area for all ranges. It encompasses many nested sub-patterns (shown as distinct colors in the previous classification).

**S2 Fig. Histograms of pairwise range overlaps among the whole community, informative taxa, and biotic elements.** When all range overlaps of the bird community are considered (first row), most have low spatial congruence similarity values (first column; mean±sd=0.13±0.18, n=1.37M overlaps), a high number of overlaps by species (second column; 1252±520, n=1095 species), and low CS means (third column; 0.12±0.06, n=1095 species). The same pattern is repeated for all overlaps of informative species (second row; n=519 species). When only overlaps composing biotic elements are considered (third row) the pattern is inverted. CS values are high (0.71±0.17; n=14597 overlaps of 558 species) for a small number of overlaps (26±34, n=558) with higher mean CS (0.61±0.19, n=558). When widespread and non-Amazonian patterns are filtered out (forth row) the pattern is modified with slightly lower means (0.57±0.2, n=5088), overlaps (11.1±12.8, n=457), and CS (0.57±0.18,n=457).

**S3 Fig. Exploratory analyses of the relationships between range overlaps, spatial congruence, and biotic elements among lowland birds of Amazonia sensu latissimo** [36]. From (B) to (J) two disproportionately large patterns (one gathering 99 transcontinental, and another 92 widespread taxa all across Amazonia) were filtered out. In general, taxa with larger ranges have higher CS means (A, r2=0.66, F1,1093=2099, p<0.01), even for biotic elements (B, r2=0.5, F1,388=392,p<0.001). However, patterns derived from larger ranges are not richer (C, r2=0). The richness of a biotic complex is not related to the mean CS (D, r2=0), and only negligible effects from spam (E, r2=0.05, F1,455=26, p<0.001) or maximum threshold values (CT) were detected (F, r2=0.11, F1,455=56, p<0.001). For all biotic complexes, CT has a negligible positive effect over the number of species and mean depth (not shown, r2=0). However, if reference species are modeled as random effects, lower CT allow more species depth and species (G, Fixed effect: accumulated number of species Ct, r2=0.84, F1,4473=2780, p<0.001; Random effect: reference species, Var=157.9±12.1, Res = 15.8, Var ratio=9.98, 90% explained), and mean depth (H, Fixed effect: mean depth, r2=0.94, F1,4473=472, p<0.001; Random effect: reference species, Var=1.89±0.13, Res = 0.11, Var ratio=17, 94.5% explained). Depth is positively related with species richness (I; r2=0.53, F1,4493=5035, p<0.001).

**S4 Fig. Species sample size, grain, and cluster settings in Infomap bioregionalizations.** Large samples have a greater chance to have species carrying contradictory biogeographic information. (A-C) With the larger sample of 1050 birds with more than 50% of its distribution in Amazonia sensu latissimo, distinct spatial scale and cluster size settings (y-axis) are not enough to recover a fine lowland spatial structure. (D-F) Within a set of species restricted (>90%) to Amazon, spatial scale adaptive resolution and cluster size adjusts show improved bioregions outlines (G-I). The use of only non-widespread informative species adds stability to the classifications across scale and depth settings.

**S5 Fig. Infomap cell-based network relationships.** Each cell has its affinity to other cells mediated by relationships with nearer cells in comparison to similarity indexes (Vilhena & Antonelli 2015). This affinity is described regionally by a set of ‘common species’, but also by ‘indicative species’ of the local composition (cell). Reference cells shown are darker blue with a red border. In complex biogeographic regions, such as Guiana, independent patterns are overlaid at biogeographic scales. This makes contiguous cells behave in an unpredictable way. A) and B) Two cells with top indicative species representing a Rio Branco river varzea specialists pattern (*Cercomacra carbonaria* and *Synallaxis collari*; [39]) mixed with Guiana [21], and Imeri [40] patterns. C) Imeri and Guiana pattern species. D) Pantepui species (*Herpsilochmus roraimae*, *Neomorphus rufipennis*) grouped with typical lowland species. E) and F) typical Guiana and Imeri species giving rise to very distinct regional patterns from contiguous cells. Patterns selected from the 329 selected species input data at a one by one degree cell (Figure S4H).

**S1 Table. Biogeographic elements in the simulated hypothetical gradient.** Among the 30 ‘species’ analyzed as references, the algorithm recognized 27 as biogeographically informative. Although smaller, the maximum depth setting of 3 (Max-depth) allowed the recognition of highly congruent patterns, as shown by the maximum and minimum congruence thresholds. These elements are enough to classify the gradient into 5 distinct non-overlapping zones (**Fig 3A**), which gain more species and expand at lower congruences (**Fig 3B)**. More relaxed depth settings allow larger groups at intermediate congruences with larger chains of indirect connections. All 23 unique biogeographic elements recovered are nested to one of the patterns grouping all species of the South or North (11 and 12 patterns, respectively; **Fig 3D**). Patterns depicted in Fig 3 are referenced in the last column.

**S2 Table. Simulated hypothetical gradient of ranges by maximum depth settings.** All possible biogeographic elements for each reference species at each distinct max-depth configuration (3, 5, 7, and 10). Number of species, spatial congruence similarity, and max. and min. values of threshold and depth for each biogeographic element are presented.

**S3 Table. Biogeographic complexes derived from selected illustrative South American bird species.** The congruence algorithm groups species with direct (depth 1) and indirect (depth > 1) relationships to the reference at each congruence threshold (C_T_). C_S_ index shows the lower C_T_ which allows a direct relationship between the grouped species and the reference. C_T_ max and min show the range of C_T_ values in which the Group species compose the biotic complex. For example, the *Icterus nigrogularis* complex at C_T_=0.73 (C_T_ max) (follow C_T_ max and depth values at Fig 4A) has *Tyrannus dominicensis* directly related (C_S_=0.73); this species links the reference to *Hydropsalis cayennensis* (depth=2), which has a direct congruence to the reference of only C_S_=0.71 (lower than the current threshold). At C_T_ max=0.42 the last species joins the group (*Inezia caudata*). Taxonomy and distribution follow BirdLife and HBW (2016).

**Table S4. Bird’s synonym and nested biogeographic complexes.** Species sharing biogeographical properties converge to synonym complexes, in which parameters, and spatial and taxonomic composition are mostly alike. Nested complexes are taxonomic subsets of larger pattern. The patterns shown at Figs 4 and 5 (references species in bold) can be generated by other species grouped at their respective biogeographic complexes. Usually the higher CS mean to all other members gives the name to the pattern. The only exception is *Heliodoxa xanthogonys*, which here, for convenience, gives its name to the ‘Tepuis1’ pattern. This is the only presented complex with nested subsets (1.1, 1.1.1). For patterns at the Amazonian margins, not all species composing the groups (S1 Table) matched the criteria of inclusion, and were not analyzed as references (e.g, some *Heliodoxa* spp. complexes).

**S1 Script. A brief tutorial to apply the congruence framework to the theoretical simulated gradient of ranges of Kreft & Jetz (2013) [34].** The tutorial and the source script are better viewed in a code editor, such as Rstudio (www.rstudio.org). The functions presented at the Tutorial S2.1 are running and integrated, but the whole framework is still in early stages of code development. Some functions are auxiliary tools, such as coherence_to_sp, which uses congruence and depth to plot range relationships in a customized way (indicating ‘internal’, ‘external’, and other spatial relations). Many others are intermediary tools called by higher hierarchical functions. New versions will maintain the functionality but the code will be refined, re-organized as a package, and uploaded opportunistically at https://github.com/cassianogatto/congruence_source (current is congruence_source_1.1.R).

**S2 Script. Source code to the congruence framework functions.** This script is meant to be maintaned and updated at https://github.com/cassianogatto/congruence_source. To use this version, save the following script (code lines) as a ‘source_congruence_1.1.R’ document in the main directory of your analyzes, and ‘source’ it using the following code: “source (“C:/my_directory/.../source_congruence_1.1.R’)”; to edit the script load it on an editor, such as RStudio https://rstudio.com/products/rstudio/).

